# A novel antiviral formulation inhibits SARS-CoV-2 infection of human bronchial epithelium

**DOI:** 10.1101/2022.10.05.510928

**Authors:** Kevin Purves, Ruth Haverty, Tiina O’Neill, David Folan, Sophie O’Reilly, Alan W. Baird, Dimitri Scholz, Patrick W Mallon, Virginie Gautier, Michael Folan, Nicola F Fletcher

## Abstract

A novel proprietary formulation, ViruSAL, has previously been demonstrated to inhibit diverse enveloped viral infections *in vitro* and *in vivo*. We evaluated the ability of ViruSAL to inhibit SARS-CoV-2 infectivity, using physiologically relevant models of the human bronchial epithelium, to model early infection of the upper respiratory tract. ViruSAL potently inhibited SARS-CoV-2 infection of human bronchial epithelial cells cultured as an air-liquid interface (ALI) model, in a concentration- and time-dependent manner. Viral infection was completely inhibited when ViruSAL was added to bronchial airway models prior to infection. Importantly, ViruSAL also inhibited viral infection when added to ALI models post-infection. No evidence of *in vitro* cellular toxicity was detected in ViruSAL treated cells at concentrations that completely abrogated viral infectivity. Moreover, intranasal instillation of ViruSAL to a rat model did not result in any toxicity or pathological changes. Together these findings highlight the potential for ViruSAL as a novel and potent antiviral for use within clinical and prophylactic settings.

## 1. Introduction

Severe acute respiratory syndrome coronavirus-2 (SARS-CoV-2), the coronavirus responsible for the current COVID-19 pandemic, primarily causes respiratory disease but is associated with a range of clinical symptoms in infected individuals (who.int). Following exposure, SARS-CoV-2 initially replicates in the human upper airway, and ciliated bronchial epithelial cells are a major target for viral replication in the early stages of infection^1^. These cells have the highest expression levels of the viral receptor, angiotensin converting enzyme-2 (ACE-2), which is concentrated on cilia^2^, and the viral entry-associated protease, transmembrane serine protease 2 (TMPRSS2)^3^. Recent studies have highlighted that variants of concern (VOC), particularly omicron, have an early replicative advantage in ciliated epithelium of the upper respiratory tract^4^, with omicron replicating more rapidly, compared with delta or wild-type virus, in primary human nasal epithelial cells^5^.

While SARS-CoV-2 initially infects the epithelium of the human upper airway and oropharynx, infection of the lower respiratory tract may occur. In a proportion of infected individuals, interstitial pneumonia develops with viral replication in type II pneumocytes, and this is an important cause of morbidity and mortality in patients with severe COVID-19 disease^6^. Intranasally delivered antivirals, either for prophylactic use, or to reduce viral shedding in infected individuals, have great potential as adjunct therapeutics in addition to vaccination and parenterally or orally administered antivirals. Therefore, development of agents that inhibit viral infection of ciliated epithelium in the upper airway should be prioritised^7^.

We previously characterised the antiviral effect of a specifically formulated emulsion of free fatty acids, ViruSAL, on infectivity of enveloped viruses *in vitro* and *in vivo*^*8*^. In the present study, we evaluated the antiviral activity of ViruSAL against SARS-CoV-2, using the highly permissive cell line, VeroE6, and human bronchial epithelial cells cultured as an air-liquid interface (ALI) to model infection of the human upper airway^2^. The potential of ViruSAL to interfere with normal mammalian structure and function was evaluated *in vitro* and *in vivo* using intranasal instillation of rats.

## 2. Results

### 2.1. ViruSAL inhibits SARS-CoV-2 infection of VeroE6 cells in a concentration dependent manner

To test whether ViruSAL inhibits SARS-CoV-2 infection following short treatment times, SARS-CoV-2 (2019-nCoV/Italy-INMI1) was incubated with ViruSAL at concentrations ranging from 3% to 0.5% for 2 minutes, or a buffer at pH 5 for 2 minutes, to control for the pH of ViruSAL, pH restored to 7 and titrated on VeroE6 cells. Infectivity was determined 48h post exposure using 50% tissue culture infectious dose (TCID50) assays or qRT-PCR to detect SARS-CoV-2 N gene (**Table 1**; Centers for Disease Control, 2020. https://www.cdc.gov/coronavirus/2019-ncov/lab/rt-pcr-panel-primer-probes.html). A concentration-dependent reduction in infectivity was observed (**Figure 1A**), and 3% ViruSAL reduced infection to undetectable levels. Confocal imaging of SARS-CoV-2 infected VeroE6 cells revealed a concomitant reduction in SARS-CoV-2 spike protein expression, in addition to dsRNA as an independent visualisation of viral replication, in cells exposed to selected concentrations of ViruSAL (**Figure 1B**). Transmission electron microscopy (TEM) of SARS-CoV-2-infected cells revealed multiple virus-like particles within single membrane vesicles in untreated and control-treated cells, consistent with endoplasmic reticulum Golgi intermediate compartments (ERGICs)^9^, as well as single virions within the cytoplasm of infected cells that were not associated with cellular organelles (**Figure 2A**). In contrast, in cells infected with SARS-CoV-2 treated with 3% ViruSAL for 2 minutes, no evidence of assembly of virions within ERGICs was observed, although individual virions were observed within the cytoplasm of VeroE6 cells (**Figure 2B**), likely representing residual input virions.

**Table 1.**
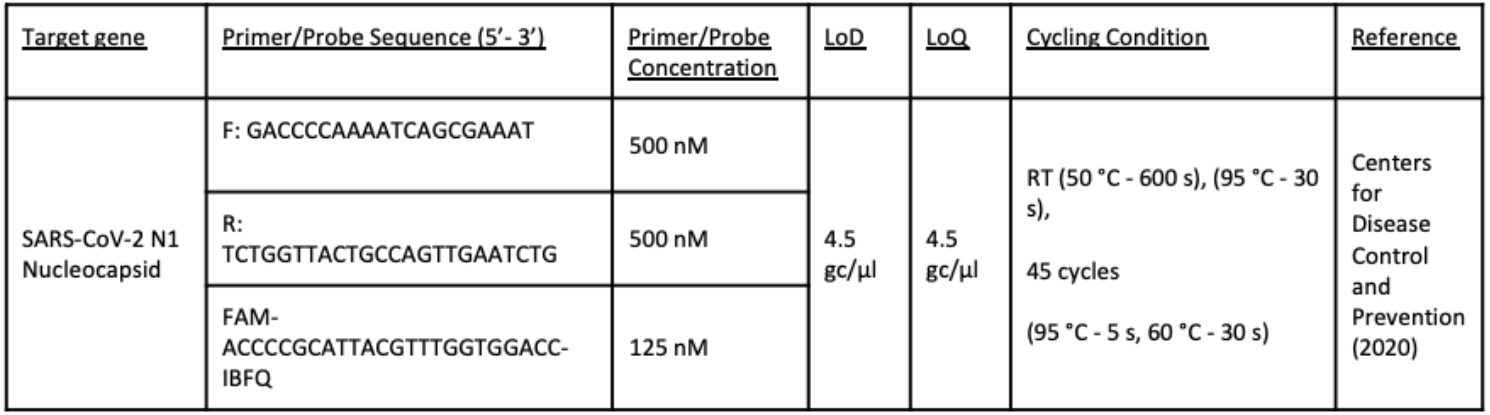

**Figure 1.**
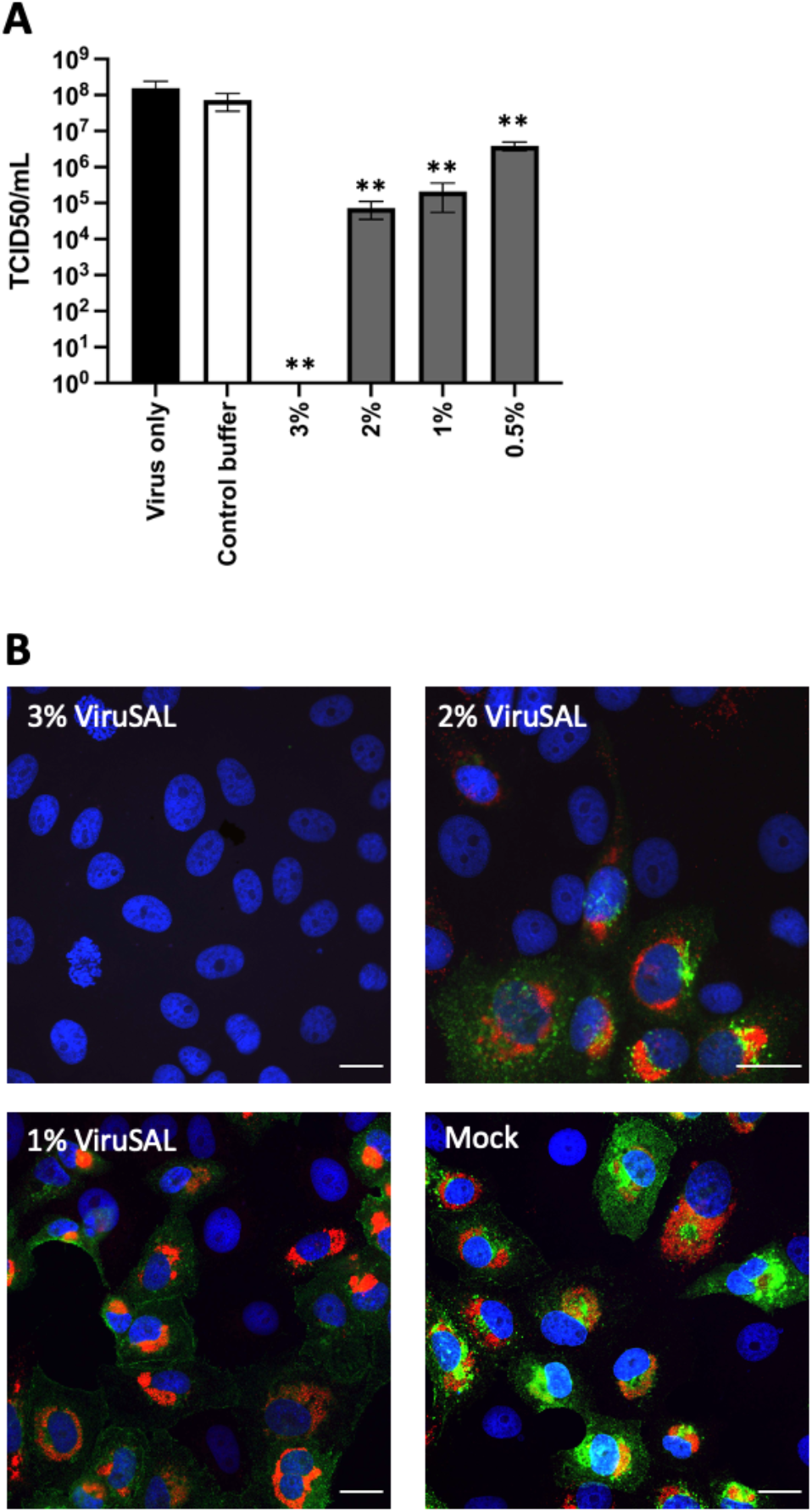
ViruSAL inhibits SARS-CoV-2 infection of VeroE6 cells in a concentration-dependent manner. SARS-CoV-2 (2019-nCoV/Italy/INMI1), at a MOI of 0.1, was treated with ViruSAL for 2 minutes, the pH restored to 7.0, and used to infect VeroE6 cells. Control cells were treated with an equal volume of pH5 buffer for 2 minutes and the pH restored to 7.0. After 48 hours, the 50% tissue culture infectious dose (TCID^50^) was calculated according to the method of Reed & Muench (1938) **(A)**. Alternatively, after 24 hours, cells were fixed and stained for SARS-CoV-2 spike protein (green), dsRNA (red) and cellular DNA (DAPI; blue) and imaged using an Olympus FV3000 confocal microscope with a 60X/NA1.4 oil lens **(B)**. Data are presented as mean infectivity ± standard deviation relative to the pH5 buffer control (n=3 independent experiments). \*\**P<0*.*01*.

**Figure 2.**
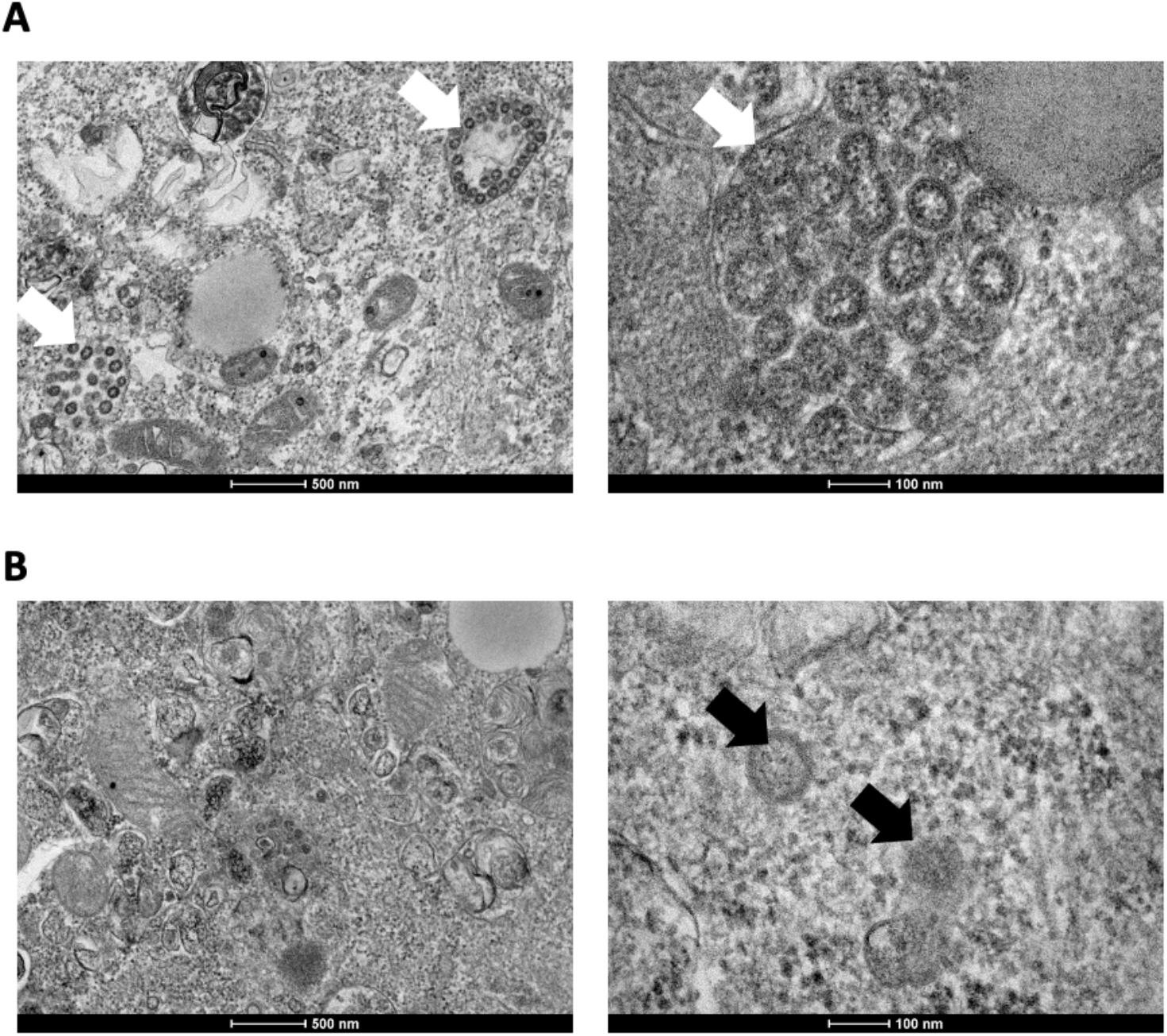
ViruSAL inhibits assembly of SARS-CoV-2 virions within VeroE6 cells. SARS-CoV-2 (2019-nCoV/Italy/INMI1), at a MOI of 0.1, either **(A)** treated with a pH5 vehicle buffer for 2 minutes as a negative control or **(B)** treated with 3% ViruSAL for 2 minutes, after which the pH was restored to 7.0 and inoculated on VeroE6 cells. After 24 hours, cells were fixed and processed for transmission electron microscopy. In control treated cells, virus particles were visible within single membrane vesicles (white arrows), whereas in ViruSAL treated cells, occasional single virions were visible within the cellular cytoplasm (black arrows).

### 2.2. ViruSAL inhibits SARS-CoV-2 of human bronchial epithelial cells cultured as an air-liquid interface (ALI)

Using human bronchial epithelial cells cultured on porous filters as an ALI culture, we determined whether ViruSAL pre-treatment of ALI cultures inhibited SARS-CoV-2 infection of the apical, ciliated surface of human bronchial epithelium. SARS-CoV-2 infection of ALI cultures was significantly inhibited by various concentrations of ViruSAL for timepoints ranging from 2-15 minutes. 3% ViruSAL reduced infectivity to undetectable levels for all timepoints measured, and a time- and concentration-dependent decrease in infectivity was observed for 2% and 1% ViruSAL (**Figure 3A,B**). These data confirm that 3% ViruSAL completely inhibits SARS-CoV-2 infection, at contact times of as low as 2 minutes, of ALI cultures prior to infection with SARS-CoV-2 via the apical surface, mimicking exposure of the ciliated bronchial epithelium of the upper airway during initial infection.

**Figure 3.**
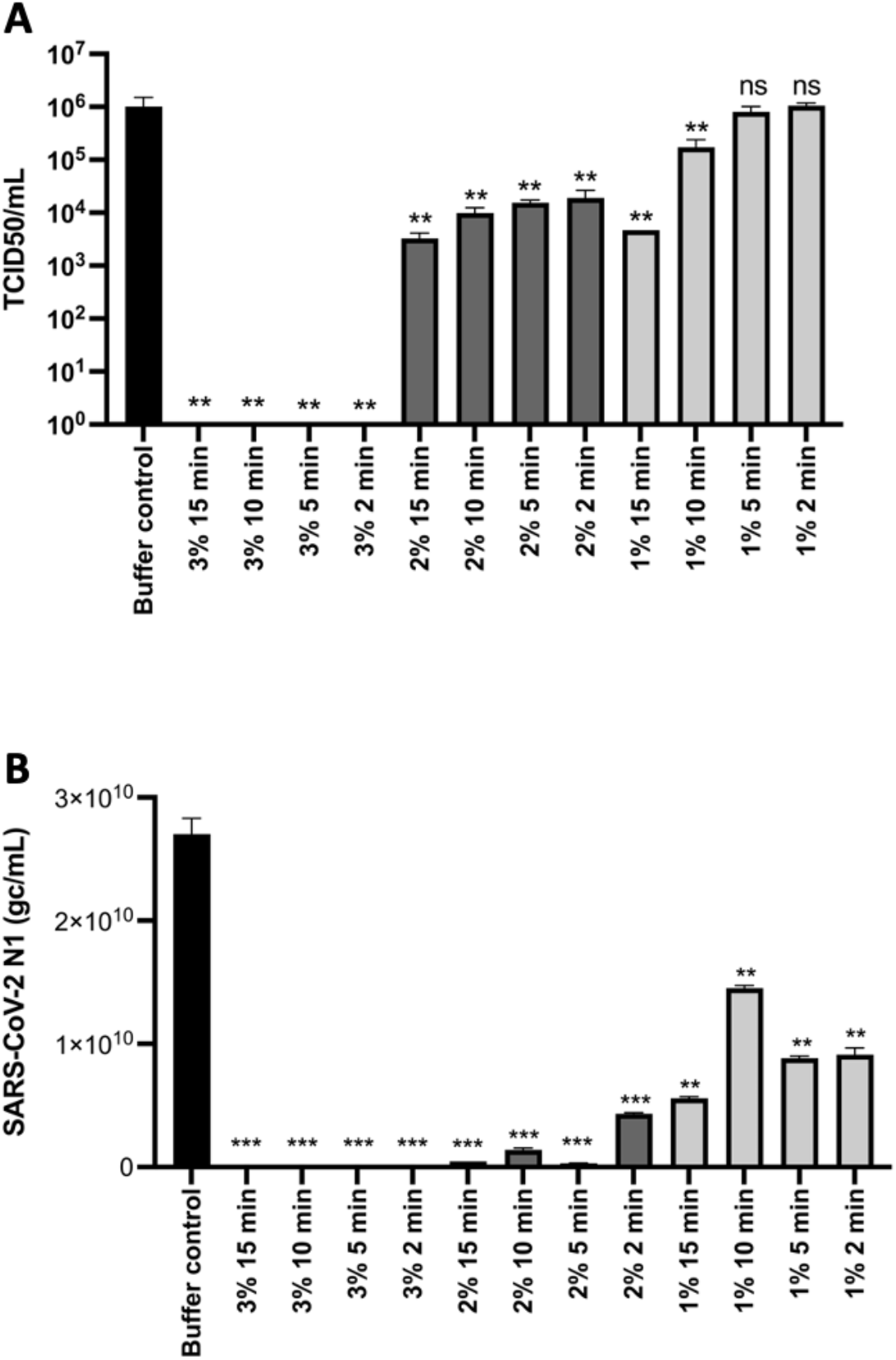
ViruSAL inhibits SARS-CoV-2 infection of human bronchial epithelial cells, cultured as an air-liquid interface (ALI), in a concentration and time-dependent manner. ALI models of human bronchial epithelial cells were treated with ViruSAL at the indicated times and concentrations, and then infected with SARS-CoV-2 (2019-nCoV/Italy/INMI1) at a MOI of 0.1 for 1 hour. After 72 hours, cell lysates were titrated on VeroE6 cells and the 50% tissue culture infectious dose (TCID50) calculated according to the method of Reed & Muench (1938) **(A)** or SARS-CoV-2 N1 quantified by qRT-PCR **(B)**. Data are presented as mean infectivity ± standard deviation relative to the pH5 buffer control (n=3 independent experiments). \*\*\**P<0*.*001*, \*\**P<0*.*01, ns: not significant*.

### 2.3. ViruSAL inhibits SARS-CoV-2 infection of ALI cultures both pre- and post-infection

Since ViruSAL potently inhibits infection of ALI cultures pre-incubation, we sought to establish whether this effect was also observed post-infection. ALI cultures were exposed to SARS-CoV-2 for 1 hour, unbound virus was removed and 3% ViruSAL was added to the apical side of ALI cultures for 15 minutes. Similar to pre-incubation with ViruSAL, exposure of ALI cultures to ViruSAL 1 hour post inoculation with SARS-CoV-2 resulted in complete neutralisation of infection, occurring in a concentration-dependent manner (**Figure 4**).

**Figure 4.**
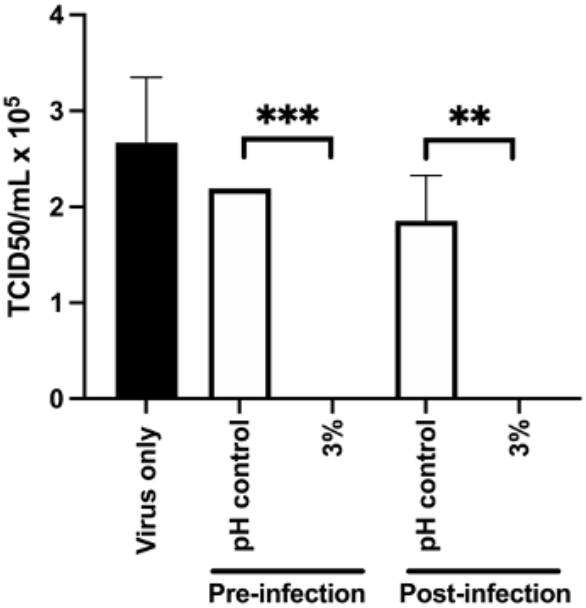
ViruSAL inhibits SARS-CoV-2 infection of ALI models pre- and post-infection. The human bronchial epithelial cell line (16HBE) was cultured as an air-liquid interface on permeable filters. Cells were treated with 3% ViruSAL for 15 minutes and then infected with SARS-CoV-2 (Strain 2019-nCoV/Italy/INIMI1) for 1 hour pre-exposure. Alternatively, cells were infected with SARS-CoV-2 for 1 hour and then treated with 3% ViruSAL for 15 minutes post-exposure. After 72 hours, cells were lysed and supernatants titrated on VeroE6 cells. 72 hours later, VeroE6 cells were scored for cytopathic effect and the 50% tissue culture infectious dose (TCID_50_) calculated. Data are presented as mean infectivity ± standard deviation relative to the pH5 buffer control (n=3 independent experiments). \*\*\**P<0*.*001*, \*\**P<0*.*01*.

### 2.4. ViruSAL inhibits SARS-CoV-2 variants of concern (VOC)

While ViruSAL inhibited wild-type SARS-CoV-2, we sought to determine whether ViruSAL can also inhibit clinical isolates representing VOC. Clinical isolates representing alpha, delta and omicron VOC were incubated with 3% or 2% ViruSAL for 2 minutes, or a pH 5 control buffer, pH restored to 7 and titrated on VeroE6 cells expressing TMPRSS2 for 48h. Infectivity was measured 48h post infection using 50% tissue culture infectious dose (TCID50) assays. 2 minute treatments with 3% or 2% ViruSAL resulted in a significant inhibition of all VOC, with 3% ViruSAL reducing alpha and omicron infectivity to undetectable levels and minimal residual infectivity of delta (**Figure 5**).

**Figure 5.**
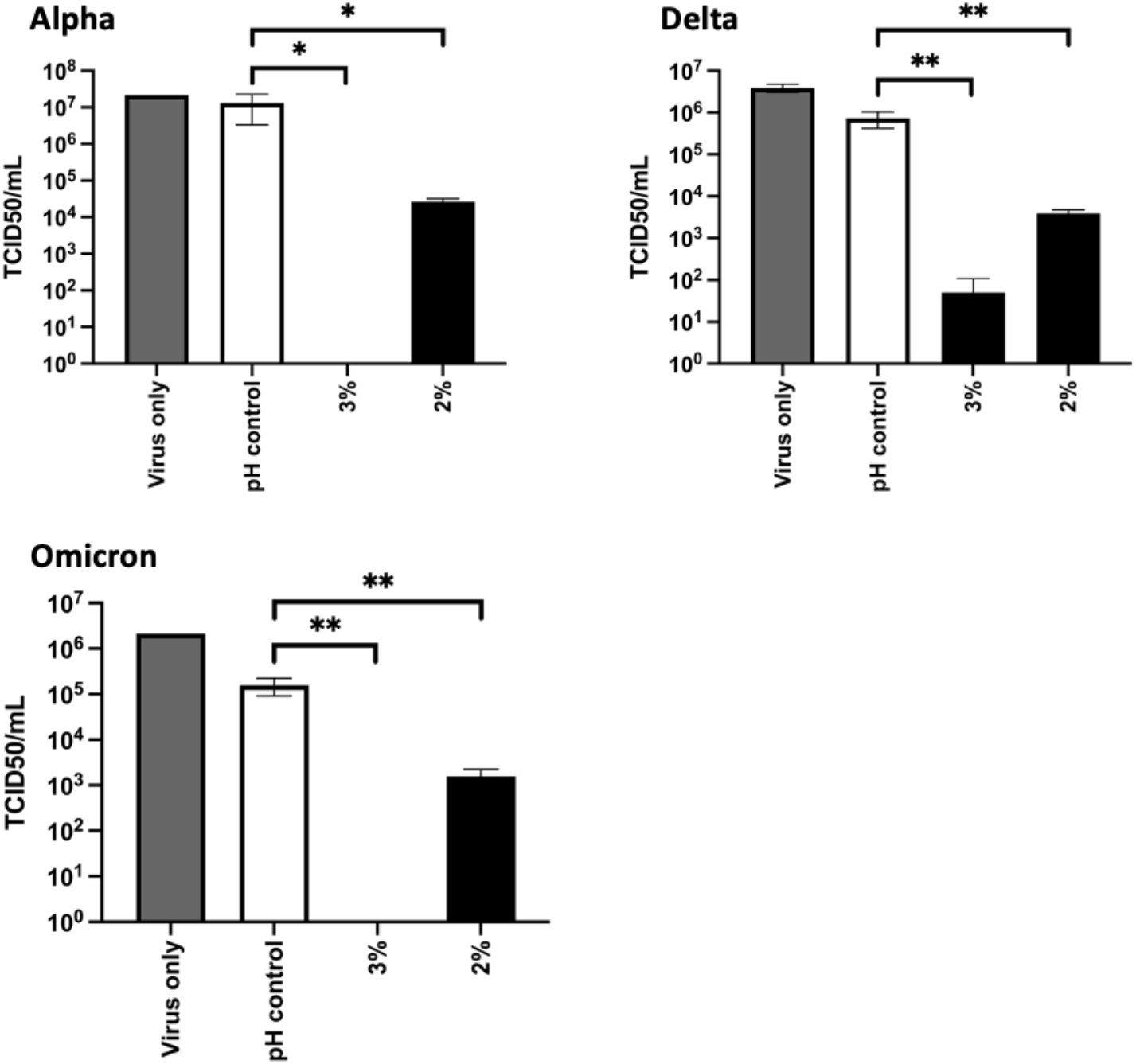
ViruSAL inhibits SARS-CoV-2 alpha, delta and omicron variants. SARS-CoV-2 clinical isolates representing alpha, delta and omicron variants, at a MOI of 0.1, were treated with ViruSAL at 3% or 2% for 2 minutes, the pH restored to 7.0, and used to infect VeroE6-TMPRSS2 cells. Control cells were treated with an equal volume of pH5 buffer for 2 minutes and the pH restored to 7.0. After 48 hours, the 50% tissue culture infectious dose (TCID_50_) was calculated according to the method of Reed & Muench (1938). Data are presented as mean infectivity ± standard deviation relative to the pH5 buffer control (n=3 independent experiments). \*\*\**P<0*.*001*, \*\**P<0*.*01*.

### 2.5. Intranasal instillation of ViruSAL does not result in clinical effects or pathological changes in rats

Twelve Sprague Dawley rats were divided into two groups of six (three males and three females per group). Group 1 was administered 25µL ViruSAL per nostril intranasally once daily for 5 days, and Group 2 received the same dosing regimen except that a pH 5 buffer was administered instead of ViruSAL. Mortality was unaffected by ViruSAL instillation and there were no premature decedents. There were no clinical signs reported for any animal at any time, and no adverse effects of dosing were evident. All animals recorded normal bodyweight profiles throughout the study, and normal food and water consumption was recorded for all animals throughout the study. There were no gross pathological changes that could be attributed to ViruSAL administration (**Figure 6A**). One animal (Female 2502 in Group 2) had red foci on all lung lobes, and they correlated histologically with haemorrhage in the alveoli that was attributed to agonal changes during euthanasia (**Figure 6B**). Three animals, one from the control group and two from the test group, had a bilateral dilation of the renal pelvis, indicative of the early stages of chronic nephropathy, which is a common spontaneous lesion in rats^10^. The lungs from each animal were weighed immediately following euthanasia, and weights were not significantly different in control and test groups (data not shown).

**Figure 6.**
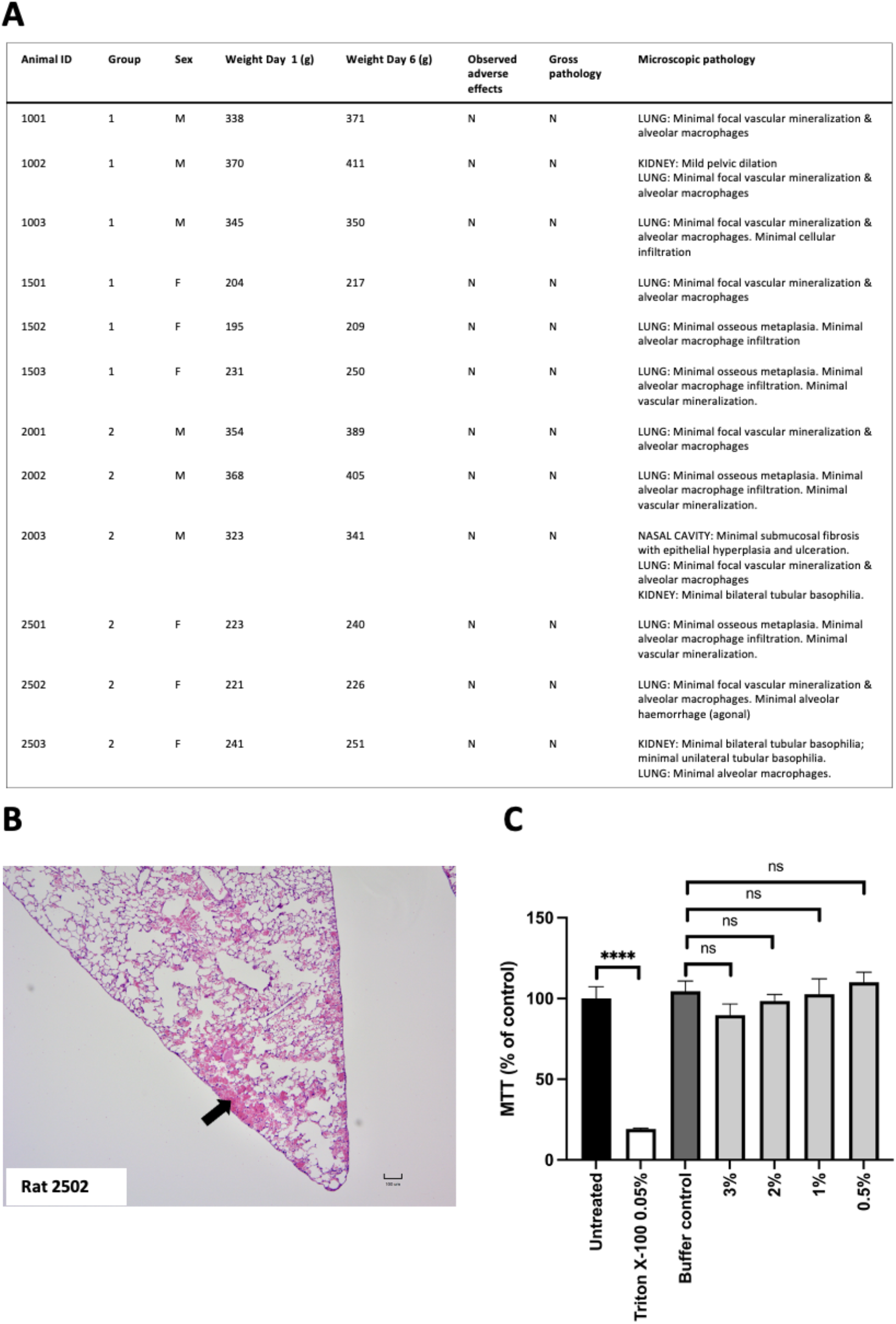
ViruSAL does not cause cellular toxicity *in vivo* or *in vitro*. The effect of ViruSAL when delivered via intranasal instillation of ViruSAL to rats for 5 days was assessed using clinical evaluations, gross and histological examination. No toxic effects of twice daily instillation of ViruSAL to the nasal cavity of rats were observed compared to a pH5 buffer control **(A)**. Mild haemorrhage was observed in the lung tissue of one rat that received ViruSAL (arrow), but this was attributed by the examining pathologist to agonal breathing during euthanasia and was unrelated to the administration of ViruSAL **(B)**. Human bronchial epithelial cells cultured as an ALI model were treated with ViruSAL, or a pH 5.0 buffer, for 15 minutes at various concentrations, and compared to 0.05% Triton-X-100 as a positive control for cellular toxicity. After 72 hours, MTT was assayed **(C)**. Data are presented as mean infectivity ± standard deviation (n=3 independent experiments). \*\*\*\**P<0*.*0001*.

Histopathologically, one animal from Group 2 (Male 2003) had a focus of minimal submucosal fibrosis with minimal hyperplasia of the overlying epithelium in the nasopalatine duct of the nasal cavity. This was considered by the pathologist to be too chronic a lesion to have developed within the five-day dosing period. This animal also had a minimal focus of ulceration with mixed cell inflammatory infiltrate in the gingiva (data not shown). Due to the absence of any abnormality in the nasal cavity in the other five animals treated with ViruSAL, the findings in the nasal cavity of this animal were considered unrelated to the administration of ViruSAL. There were no other clinical signs, gross pathological changes or histopathology which could be related to ViruSAL administration **(Figure 6A,B)**.

### 2.6. ViruSAL treatment of ALI cultures results in minimal disruption of cellular viability

We previously demonstrated that ViruSAL treatment does not disrupt cellular viability of several cell lines and primary cell cultures^8^. To investigate whether ViruSAL affects viability of human ALI cultures, the apical surface of models were treated with ViruSAL, or pH 5.0 control buffer, for various timepoints ranging from 2 to 15 minutes, at concentrations ranging from 3% to 0.5%, and viability measured by MTT assay. Cellular viability was not significantly reduced compared with pH 5 treated buffer in cells treated with ViruSAL for various concentrations up to 3% ViruSAL for 15 minutes (**Figure 6C**).

## 3. Discussion

The socioeconomic and public health impacts of the ongoing COVID-19 pandemic have highlighted the need for the development of broad-spectrum antivirals for this and future viral outbreaks^11^. Moreover, despite the availability of antivirals and vaccines targeting SARS-CoV-2, intranasally delivered antivirals, with prophylactic applications or to reduce viral shedding in infected individuals, have significant value as adjunct therapeutics particularly in healthcare settings and other environments with high risk of viral transmission^7^.

Following infection, SARS-CoV-2 initially replicates in the nasopharyngeal epithelium of the upper respiratory tract, where it attaches to target cells initially via binding to the ACE-2 receptor that is expressed on cilia on the apical surface of the cells^1,6^. Following viral replication and assembly, viral particles are shed from infected cells where they may be transmitted to another host via close contact including coughing. Furthermore, in a proportion of individuals, SARS-CoV-2 infects the lower respiratory tract or systemic sites including the kidney, pancreas, heart and also the central nervous system^12^. Therefore, the upper respiratory tract is an important site for initial exposure, infection, viral transmission and dissemination. The air-liquid interface of bronchial epithelial cells cultured on porous inserts represents a physiologically relevant system to investigate initial SARS-CoV-2 infection and to evaluate antivirals targeting this site. Epithelial cells cultured as ALI models develop polarity that is similar to the *in vivo* state, with cilia development on the apical (air) side that express ACE-2 exclusively and have been used to model respiratory virus infections including SARS-CoV and SARS-CoV-2^13,14^. ALI models are a relevant model system to evaluate antiviral agents against SARS-CoV-2, and remdesivir and other antivirals have been evaluated using this model^15,16^.

We previously reported that ViruSAL is a specifically formulated emulsion of free fatty acids which has antiviral properties against a range of enveloped viruses *in vitro* and *in vivo*^*8*^. *In vitro*, ViruSAL potently inhibited replication of a range of enveloped viruses, including Epstein-Barr, measles, herpes simplex, Zika and orf parapoxvirus, together with Ebola, Lassa, vesicular stomatitis and severe acute respiratory syndrome coronavirus (SARS-CoV) pseudoviruses. *In vivo*, ViruSAL significantly inhibited Zika and Semliki Forest virus replication in mice following the inoculation of these viruses into mosquito bite sites. Caprylic acid is the major antiviral component in ViruSAL, however its emulsification within the ViruSAL formulation enhances its antiviral properties and stability compared to caprylic acid alone. Transmission electron microscopy analysis revealed that ViruSAL disrupted the integrity of parapox virions, with disruption of the viral envelope. In addition, ViruSAL had no effect on non-enveloped viruses, indicating that its mechanism of action may involve inhibition of viral entry through disruption of the viral envelope^8^. This is a novel approach to disrupt the phospholipid membranes of diverse enveloped viruses. In this study, we report that ViruSAL has antiviral activity against wild type, and variants of, SARS-CoV-2, in both the highly permissive VeroE6 cell line and an ALI model of human bronchial epithelium. Therefore, these data align with previous findings, which indicated that ViruSAL significantly decreased infectivity across a range of enveloped viruses (Zika, herpes simplex, measles, Epstein-Barr and orf parapoxvirus)^8^.

ViruSAL also inhibited viral infectivity even when applied to ALI cultures post-infection. The mechanism by which ViruSAL inhibits viral infectivity post-infection remains to be elucidated. Our previous reports demonstrate that ViruSAL reduces Zika and Semliki Forest virus local replication of mouse skin when applied post viral challenge, as well as viraemia and dissemination to spleen^8^. Previous studies investigating the mechanism of action of monocaprylate, a monoglyceride homologue of caprylic acid, have also highlighted destabilisation of bacterial membrane by increasing permeability and the number of phase boundary defects^17^.

Multiple reports have demonstrated that coronaviruses including SARS-CoV-2, which replicate within the cytoplasm and assemble at intracellular membranes. During replication, SARS-CoV-2 modifies cellular membranes to form double membrane vesicles (DMVs) that are associated with the endoplasmic reticulum-Golgi intermediate compartment (ERGIC), which is the main site of SARS-CoV-2 assembly^18^. Transmission electron microscopy (TEM) of infected VeroE6 cells treated with a control buffer (mock treatment) or 3% ViruSAL revealed multiple aggregations of particles consistent with SARS-CoV-2 virions within DMVs in mock treated cells, indicating viral replication and assembly within these cells. These structures were not observed within virus inoculated VeroE6 cells treated with ViruSAL, indicating that ViruSAL inhibited viral infection. In both ViruSAL treated and mock treated cultures, low numbers of individual virus particles were visible within the cytoplasm of cells, which likely represents residual input virus despite removal of inoculum 1h post infection followed by washing of the cells. The lack of virus-containing DMVs in ViruSAL-treated cells indicates that input virions were unable to replicate in treated cells, in contrast to mock treated cultures. Therefore, our data suggests that the mechanism of action of ViruSAL likely includes prevention of viral replication by disrupting the viral membranes of progeny virions, as previously reported for other enveloped viruses^8^.

We previously reported that ViruSAL exerts minimal effects on cellular viability of diverse cell lines *in vitro*^*8*^. Viability of ALI models following ViruSAL treatment was assessed to determine if there is any adverse cellular toxicity, and as previously observed, treatment of cells with ViruSAL did not significantly alter viability of ALI cultures at antiviral concentrations. Similarly, no adverse clinical signs or gross or histopathological evidence of toxicity was observed *in vivo* following intranasal instillation of ViruSAL to rats, following twice daily treatments for five days. These data highlight the potential of ViruSAL as an adjunct topical antiviral against SARS-CoV-2 infection of the upper airway.

In conclusion, the present study demonstrates that ViruSAL is a potent antiviral formulation with activity against SARS-CoV-2 both pre- and post-infection in an ALI model of the human upper airway. Antiviral activity of ViruSAL occurs in a time- and concentration-dependent manner. Following ViruSAL treatment no evidence of viral replication was observed in the highly permissive cell line, VeroE6. In addition, no evidence of cellular toxicity was observed *in vitro* or *in vivo* when delivered intranasally to rats. Taken together, this study demonstrates the potential of ViruSAL as a preventive or adjunct therapy for SARS-CoV-2 and other enveloped respiratory viruses^8^. ViruSAL is potentially a valuable addition to the limited array of broad-spectrum antivirals for infections of mucosal surfaces and the respiratory tract.

## 4. Materials and methods

### 4.1. Cell lines and antibodies

VeroE6 cells (ATCC) and VeroE6 cells transiently expressing an untagged TMPRSS2 cDNA expression vector (Sinobiological; HG13070-UT) were propagated in DMEM supplemented with 10% fetal bovine serum (FBS), 2 mM L-glutamine (Gibco; Thermo Fisher, MA, USA), and 1% non-essential amino acids (Gibco). Normal human bronchial epithelial cells were obtained from Lonza (Basel, Switzerland; CC-2540S) and cultured in B-ALI medium (Lonza) according to the manufacturer’s instructions. To generate ALI cultures, 1 × 10^5^ cells were seeded in the upper compartment of Transwell filter inserts (Corning, AZ, USA) with a 0.4mm pore size and cultured for 72 hours. Culture medium was then removed from the upper compartment of the inserts and cells were cultured for a further 14 days to facilitate cellular differentiation and cilia development. Antibodies used were anti-rabbit SARS-CoV-2 spike S1 antibody (Sinobiological, USA) and anti-mouse dsRNA K1 antibody (Scicons, Hungary). Secondary antibodies were anti-mouse Alexa Fluor 594 and anti-rabbit Alexa Fluor 488 antibodies (Thermo Fisher, MA, USA).

### 4.2. Virus culture and infection assays

SARS-CoV-2 (2019-nCoV/Italy-INMI1 from EVA Global) was propagated in VeroE6 cells. All virus used in this study was at passage 3. Cells were inoculated with an MOI 0.01 for 2 hours, washed with PBS, and the medium replaced with DMEM containing 2% FBS. When cultures were fully infected, flasks were freeze-thawed three times and supernatants collected and clarified at 3500 rpm for 30 minutes at 4°C. Supernatant was collected, aliquoted and stored at -80°C. TCID_50_ was performed in VeroE6 cells in quadruplicate and infectious titre determined using the Reed-Muench method^19^. SARS-CoV-2 variants including Alpha (CEPHR_IE_B.1.1.7_0221, GenBank accession ON350867, passage 2), Delta (CEPHR_IE_AY.50_0721, GenBank accession ON350967), Omicron (CEPHR_IE_BA.1_0212, GenBank accession ON350968, Passage 2) clinical isolates were isolated from SARS-CoV-2 positive nasopharyngeal swabs from the All-Ireland Infectious Disease (AIID) cohort^20^ were isolated and amplified on Vero E6/TMPRSS2 cells (#100978), obtained from the Centre For AIDS Reagents (CFAR) at the National Institute for Biological Standards and Control (NIBSC)^21^

For fluorescence microscopy and transmission electron microscopy, cells were infected for 16-24 hours with SARS-CoV-2 at an MOI of 0.01 prior to fixation and processing.

### 4.3. Confocal and transmission electron microscopy

VeroE6 cells infected with SARS-CoV-2 for 24 hours as described in Section 4.2 were fixed for 1 hour with 4% paraformaldehyde at 37°C, washed with phosphate buffered saline (PBS), and then incubated with 0.5 %(v/v) Triton X-100 and 0.5 %(w/v) Bovine Serum Albumin in PBS (blocking buffer) for 30 minutes at room temperature. Primary antibodies SARS-CoV-2 Spike S1 Antibody rabbit mAb (SinoBiological, 40150-R007, USA) and anti-dsRNA K1 mAb mouse IgG2a (Scicons K1-1910, Hungary) in blocking buffer were incubated for 1 hour at room temperature. The cells were washed 5x with blocking solution and a 1:500 dilution of secondary antibodies (Invitrogen Goat anti-rabbit IgG (H+L) Superclonal™/Invitrogen Alexa Fluor™ 594 goat anti-mouse IgG2a (γ2a) in blocking solution was incubated for 1 hour at room temperature. The cells were washed 5x with blocking solution and mounted with Invitrogen ProLong™ Gold antifade reagent with DAPI (Thermo Fisher, MA, USA). Slides were imaged with an Olympus FV3000 confocal microscope.

For transmission electron microscopy (TEM), VeroE6 cells infected with SARS-CoV-2 for 24 hours as described in Section 4.2 were fixed for 1 hour with 2.5% glutaraldehyde, post fixed with 1% osmium tetroxide and dehydrated using a graded ethanol series (30%, 50%, 70%, 90%, 100%). Dehydrated samples were transferred into acetone and then embedded in Epon. Ultrathin (80 nm) sections were obtained using a Leica EM UC6. These sections were collected on 200 mesh thin bar copper grids, stained with uranyl acetate and lead citrate. Sections were visualized using a Tecnai G^2^ 12 BioTWIN transmission electron microscope.

### 4.4. Construction of ViruSAL

ViruSAL was constructed by Westgate Biomedical Ltd, Ireland, (Folan M. Patent WO 2011/061237), and provided to UCD as a gift under a research agreement. All components used in the construction of ViruSAL are pharmaceutical grade constituents of greater than 97% purity. ViruSAL was constructed as previously described^8^.

For *in vivo* safety evaluation as described below at 5.8 the co-surfactant Pluronic F68 was replaced with Tween 80 at 0.15% in the concentrate, lecithin S75 and DPPG were used at 1.2% and 0.36%, respectively, and the oil phase totalling 10% was constructed using caprylic acid, Miglyol 812 and peppermint oil (8:1:1). In this case the emulsion was formed using a Niro Sauvi PandaPLUS 2000 homogeniser (GEA, Italy) with 4 passes at 900 bar. The emulsion concentrate was diluted to 3% w/w in 50 mM sodium citrate buffer at pH 4.7 for use in the intranasal instillation study and a control formulation consisting of 50 mM sodium citrate buffer at pH 4.7 was also used in that study. Caprylic acid is the functional surfactant in ViruSAL, all other constituents are excipients which facilitate delivery. The 3% w/w dilution used in the intranasal instillation study contained 0.24% w/w caprylic acid.

### 4.5. Assessment of the effect of ViruSAL on viral infection *in vitro*

When evaluating the effect of ViruSAL on SARS-CoV-2 infection of VeroE6 cells, ViruSAL at the indicated concentrations was mixed with an equal volume of viral inoculum (TCID_50_ = 4.48^7^ mL^-1^) for specified times after which the same volume of a buffered neutralizing solution was added and mixed thoroughly to restore the pH to 7.0. ViruSAL and neutralizing solution was optimized for the DMEM/2% FBS used in *in vitro* studies. As a control for the effect of low pH on viral infectivity, an equal volume of pH 5.0 buffer solution was added to virus, incubated for 2 minutes and neutralized as before. Virus/ViruSAL, or virus mock treated with pH 5.0 buffer, was inoculated onto VeroE6 cells and incubated for 72 hours at 37 °C, then infection quantified using the TCID^50^ assay.

For air-liquid interface (ALI) models, ViruSAL or pH 5.0 buffer control was added directly to the apical (‘air’) side of the models and incubated for various timepoints. SARS-CoV-2 was then added and incubated for 1 hour on the apical side of the ALI cultures, cells washed once with pre-warmed PBS and incubated for 72 hours. Alternatively, to evaluate the ability of ViruSAL to inhibit viral infection post-infection, cells were incubated with SARS-CoV-2 for 1 hour, washed with PBS and ViruSAL was added to the apical side of ALI cultures for 15 minutes and cultures incubated for 72 hours. 72 hours post infection, 200 µl culture media was added to the apical side of ALI cultures and they were freeze-thawed 3 times and centrifuged at 3,000 rpm for 15 minutes. Supernatant was titrated on VeroE6 cells, incubated at 37 °C for 72 hours and cytopathic effect (CPE) evaluated via microscopic visualization. TCID_50_ was calculated according to the method of Reed and Muench^19^.

### 4.6. MTT assay

To assess cell viability following ViruSAL treatment, an MTT assay (Vybrant MTT Cell Proliferation Assay, Thermo Fisher Scientific, U.K.) was performed on ALI cultures 72 hours following ViruSAL treatment, to mimic the timepoint at which viral infection was measured. The MTT assay was performed according to the manufacturer’s instructions and as previously described^8^.

### 4.7. RNA Extraction and RT-qPCR

RNA was extracted from cellular material using the QIAamp Viral RNA Mini kit (Qiagen, Germany) according to the manufacturer’s instructions. RT-qPCR was performed as previously described^22^. Primer/probe and reaction details are described in Table 1. All RT-qPCR assays were performed in a total volume of 20 μL, including 5 μL template using LightCycler Multiplex RNA Virus Master (Roche, Switzerland). All assays were carried out in triplicate on the Roche Lightcycler 96 platform (Roche, Switzerland) with four quantification standards included.

### 4.8. Safety evaluation of intranasal instillation of ViruSAL to rats

To evaluate any potential toxicity following administration of ViruSAL, a study evaluating the effect of intranasal instillation of ViruSAL to a rat model was carried out by Charles River Laboratories. All experiments were performed in accordance with relevant local guidelines and authorisations, and the study was carried out under Home Office Project Licence No. PBAD559F8, Toxicology of Pharmaceuticals, Protocol No. 1.

12 Sprague Dawley rats (6 male, 6 female), of at least 8 weeks of age and 200-300 g (males) or 150-200 g (females) were randomly assigned to two groups, with an equal number of males and females per group. Animals were group housed, with up to 3 animals of the same sex and dosing group housed together. Group 1 (control group) was administered 25 µL 50 mM citrate buffer in each nostril using a micropipette with an appropriate sized plastic tip once daily for five days. Group 2 (test group) was administered 25 µL 3% ViruSAL (equivalent dose of 1500 µg/animal) into each nostril once daily for five days, using the same procedure as described for the control group. Animals were observed within their cage for 1-2 hours post-dose daily and at least once daily at other times post-dose for identification of possible adverse effects. Detailed clinical observations were made and animal weight recorded twice weekly, and food and water consumption were monitored daily. Animals were euthanised on day 6 by exposure to rising levels of carbon dioxide, followed by recording of terminal body weight and then exsanguination. Animals were subjected to a complete necropsy examination and gross observations and organ weights as a percent of brain weights recorded. Representative samples of tissues were fixed in 10% neutral buffered formalin for histopathologic and microscopic examination by a veterinary pathologist trained in laboratory animal pathology.

### 4.9. Statistical analysis

*In vitro* results are expressed as the mean ± 1 standard deviation of the mean (SD), except where stated. Statistical analyses were performed using Student’s t-test or One Way ANOVA in Prism 9.0 (GraphPad, USA).

## Ethical Statement

Experiments in rats were performed by Charles River Laboratories, Co. Mayo, Ireland, in accordance with relevant local guidelines and authorisations, and the study was carried out under Home Office Project Licence No. PBAD559F8, Toxicology of Pharmaceuticals, Protocol No. 1.

## Acknowledgements

The authors acknowledge Dr John Browne, School of Agriculture, University College Dublin, Ireland and Dr Bridget Hogg and Mr Marc Farrelly, School of Veterinary Medicine, University College Dublin, Ireland, for valuable technical assistance. The authors acknowledge the management of the Containment Level 3 laboratory at the School of Veterinary Medicine, University College Dublin, Ireland, for facilitating experiments with SARS-CoV-2.

